# PDB NextGen Archive: Centralising Access to Integrated Annotations and Enriched Structural Information by the Worldwide Protein Data Bank

**DOI:** 10.1101/2023.10.24.563739

**Authors:** Preeti Choudhary, Zukang Feng, John Berrisford, Henry Chao, Yasuyo Ikegawa, Ezra Peisach, Dennis W. Piehl, James Smith, Ahsan Tanweer, Mihaly Varadi, John D. Westbrook, Jasmine Y. Young, Ardan Patwardhan, Kyle L. Morris, Jeffrey C. Hoch, Genji Kurisu, Sameer Velankar, Stephen K. Burley

**Author notes:** Author notes. Contributed equally to the work (Preeti Choudhary, Zukang Feng). deceased (John D. Westbrook).

## Abstract

The Protein Data Bank (PDB) archive is the global repository for public-domain 3D biomolecular structural information. The archival nature of the PDB presents certain challenges pertaining to updating or adding associated annotations from trusted external biodata resources. While each Worldwide PDB (wwPDB) partner has made best efforts to provide up-to-date external annotations, accessing and integrating information from disparate wwPDB data centers can be an involved process. To address this issue, the wwPDB has established the PDB Next Generation or NextGen Archive, developed to centralize and streamline access to enriched structural annotations from wwPDB partners and trusted external sources. At present, the archive provides mappings between experimentally determined 3D structures of proteins and UniProt amino acid sequences, together with domain annotations from Pfam, SCOP2, and CATH databases, and intra-molecular connectivity information. Since launch, the PDB NextGen Archive has seen substantial user engagement with over 3.5 million data file downloads, ensuring researchers have access to accurate, up-to-date, and easily accessible structural annotations.

**Database URL:** http://www.wwpdb.org/ftp/pdb-nextgen-archive-site

## Introduction

The Protein Data Bank (PDB) has a remarkable history, which began in 1971 when it was established as the first open-access digital data resource in biology(1). After more than five decades of continuous operations, the PDB has grown more than 30,000 fold, increasing from seven to more than 210,000 structures and becoming a global repository storing extensively annotated 3D structures of proteins and nucleic acids, enabling atomic-level insights into the workings of complex biological macromolecules(2). This invaluable public-domain three-dimensional (3D) structure data resource has had a profound impact on fundamental biology, biomedicine, biotechnology, and bioenergy, encompassing both naturally occurring and engineered biomolecules(3,4). Moreover, the PDB has significantly contributed to human health, facilitating discovery and development of nearly 90% of new drugs approved by the United States Food and Drug Administration between 2010 and 2016 with open access to 3D biostructure information(5).

Since 2003, the PDB has been managed by the Worldwide Protein Data Bank (wwPDB) partnership ensuring that all archival data are freely accessible without limitations of its usage(6). At that time, this international partnership brought together the RCSB Protein Data Bank (RCSB PDB)(7), Protein Data Bank in Europe (PDBe)(8), and Protein Data Bank Japan (PDBj)(9) as founding wwPDB members to oversee the PDB Core Archive, which houses atomic-level 3D structures of biological macromolecules. These experimentally determined structures are the products of macromolecular crystallography (MX), nuclear magnetic resonance (NMR) spectroscopy, and electron microscopy (3DEM). Over more than two decades, the wwPDB has grown to encompass two additional full members (Biological Magnetic Resonance Data Bank or BMRB(10), responsible for archiving experimental NMR spectroscopy data; and Electron Microscopy Data Bank (EMDB)(11), responsible for archiving experimental 3DEM data), and its first associate member (PDB China or PDBc)(12). The wwPDB adheres to the principles of Fairness-Accuracy-Confidentiality-Transparency (FACT)(13) and Findability-Accessibility-Interoperability-Reusability (FAIR)(14), ensuring equitable sharing and responsible management of the 3D biostructure data. Information stored in the PDB is made available under the most permissive Creative Commons CC0 1.0 Universal License(https://creativecommons.org/licenses/by/4.0/), enabling researchers around the world to access and utilize the information. Recognizing its commitment to high standards of data management, preservation, and openness, the PDB is accredited by CoreTrustSeal, an international organization that certifies data repositories (https://amt.coretrustseal.org/certificates/). More recently, the PDB was recognized by the Global Biodata Coalition (https://globalbiodata.org) as a Global Core Biodata Resource, of “fundamental importance to the wider biological and life sciences community and the long-term preservation of biological data.”

Each PDB structure (or entry) is made up of 3D atomic coordinates, experimental data, and extensive metadata, providing a wealth of information regarding sample provenance and the structure determination process. PDB metadata encompasses protein names, amino acid or nucleic acid sequences, source organism(s), small-molecule chemical information, data collection information (*e*.*g*., instrumentation), structure-determination methodology (*e*.*g*., model-building procedures, structure quality metrics). Each PDB entry is deposited, biocurated, and validated using the common global OneDep(15) deposition, biocuration, and validation tool. During validation(16), OneDep computes quantitative assessments of structure quality, including both chemical geometry and agreement with experimental data. During the biocuration(17), expert wwPDB staff members utilize OneDep(15) to incorporate sequence cross-references to trusted external biodata resources, including UniProtKB(18) or NCBI GenBank(19), which provide links to reference sequence information. Other biocuration activities support inclusion of other derived metadata, including structural characteristics (*e*.*g*., secondary and quaternary structure) and small-molecule ligand descriptors.

The PDB weekly release process collects new 3D biostructure data from each wwPDB partner, cross-checks, and packages it into a pre-release FTP area, and then delivers this information to wwPDB partners for distribution from regional wwPDB FTP servers. On average, more than 300 new structures are publicly released into the PDB Core Archive every Wednesday at 00:00 Universal Time Coordinated (UTC). The wwPDB FTP servers support open access to the entire contents of the PDB archive, with no login requirement of usage limitations. The downloadable information is archival in nature, faithfully reflecting the 3D biostructure data generously contributed by more than 60,000 structural biologists since 1971.

Metadata associated with each newly released PDB structure are largely fixed at the time of public release. Recognizing that knowledge of biochemical, cellular, and organismal context is often necessary to understand and fully utilize 3D biostructure data, the founding wwwPDB partners (RCSB PDB, PDBe, and PDBj) serve additional, regularly updated functional annotations and related metadata with each PDB structure on their respective websites (rcsb.org, pdbe.org, and pdbj.org, respectively). In parallel, BMRB (https://bmrb.io/) and EMDB (https://www.ebi.ac.uk/emdb/) offer similarly valuable, regularly updated metadata for NMR and 3DEM structures, respectively. This arrangement provides valuable services to many millions of PDB data consumers around the world. Feedback from our diverse user community, however, requested that we augment the structure data files served *via* FTP, HTTP and rsync with up-to-date functional annotations and related metadata provided on the RCSB PDB, PDBe, PDBj, BMRB, and EMDB web portals. Because much of the contextual information provided by on these wwPDB websites is complementary, the wwPDB partnership elected to pool functional annotations and other metadata attributed to each structure and develop a “Next Generation” PDB data repository, thereby providing an efficient mechanism for sharing value-added information accumulated across the wwPDB with PDB data consumers around the world.

Herein, we describe the design and implementation of the PDB NextGen Archive, as a regularly updated (or living) one-stop-shop resource that enhances the accessibility and utilization of structural information within its biological context. The wwPDB vision for the PDB NextGen Archive is to deliver up-to-date metadata and functional annotations through an integrated and standardized methodology, thereby facilitating a comprehensive grasp of the biochemical, cellular, and organismal contexts associated with protein structures. This paper presents technological advances and the collaborative efforts of the wwPDB partnership to develop the PDB NextGen Archive. It outlines data retrieval and integration mechanisms of each wwPDB partner, showcasing a dynamic, living data resource. Launch of the PDB NextGen Archive marks a significant milestone in the evolution of the PDB as a global data resource, giving researchers the wherewithal to explore protein structures with enriched biological context and current functional annotations. It underscores the commitment of wwPDB partners to meet the evolving needs of the biological and biomedical research and education communities and to support ground-breaking research across fundamental biology, biomedicine, biotechnology, and bioenergy.

## Implementation and Outcomes

Initially introduced as a successor to the legacy PDB file format, the PDB exchange/macromolecular Crystallographic Information Framework (PDBx/mmCIF)(20–22) now serves as the master format for the PDB archive, addressing the growing complexity and diversity of structural biology data(23). The PDBx/mmCIF data standard/data dictionary overcomes limitations of the legacy PDB file format and readily accommodates PDB data encompassing very large structures of macromolecular machines and viruses, complex chemistry, and new experimental methods. The benefits of the PDBx/mmCIF format are manifold. It is both human and machine-readable. The metadata framework of the dictionary specifies data content and encompasses data typing, validation rules, and organizational structures. This comprehensive approach ensures that data consistency and integrity can be maintained *via* automated checks. The PDBx/mmCIF data standard/data dictionary is fully extensible, permitting incorporation of new data items and categories as evidenced by the X-ray Free Electron Laser/Serial Crystallography (XFEL/SX)(2). Moreover, the core PDBx/mmCIF standard promotes data sharing and interchangeability as evidenced by launch of the specialized IHMCIF(24) and ModelCIF(25) dictionaries built atop the PDBx/mmCIF data standard. By facilitating inclusion of new information and accommodating scientific advances, the PDBx/mmCIF dictionary can provide enduring value to the scientific community.

Maintenance of the PDBx/mmCIF data standard/data dictionary is overseen by the wwPDB partnership, which is continuously updating it with new data elements to accommodate new science and technology. The wwPDB collaborates with the wwPDB PDBx/mmCIF Working Group, leveraging domain experts to help refine and extend the data model. This collaborative effort ensures that the PDBx/mmCIF dictionary remains relevant and adaptable to researchers’ evolving needs. Ongoing improvements are shared with the scientific community *via* the GitHub platform (https://github.com/pdbxmmcifwg), promoting transparency, collaboration, and community involvement. To streamline access and utilization of PDBx/mmCIF, a dedicated data portal site (https://mmcif.wwpdb.org/) has been established. This comprehensive resource provides access to data standards, metadata specifications, tutorials, and links to essential software tools. By offering these resources, the wwPDB ensures that researchers can effectively navigate and harness the myriad capabilities of PDBx/mmCIF.

Based from the PDBx/mmCIF format, the Protein Data Bank Markup Language (PDBML) supports representation of PDB data in XML format(26). This schema is the product of direct translation of the PDBx/mmCIF Dictionary and is available for download from https://pdbml.wwpdb.org/pdbml-downloads.html. The NextGen Archive supports both PDBx/mmCIF and PDBML formats, providing researchers with two options for accessing and utilizing PDB data.

Development of the PDB NextGen Archive prototype began with incorporation of Structure Integration with Function, Taxonomy, and Sequence (SIFTS)(27,28) annotations, a crucial step driven by previous efforts aimed at expanding the PDBx/mmCIF data dictionary(29). This extension was designed to seamlessly integrate SIFTS data, which includes annotations from UniProtKB(18), Pfam(30), SCOP2(31), and CATH(32). Value-added, residue-level annotations were directly incorporated into the PDBx/mmCIF files from the PDB Core Archive. [_pdbx_sifts_xref_unp_segments](https://mmcif.wwpdb.org/dictionaries/mmcif_pdbx_v50.dic/Categories/pdbx_sifts_unp_segments.html) and _pdbx_sifts_xref_db_segments data categories offer information on PDB segments mapped to UniProt and other external databases (Pfam, SCOP2, and CATH), while _pdbx_sifts_xref_db provides a complete view of all residue mappings to these trusted external resources (Figure 1). The best-mapped UniProt residue numbering was also incorporated in _atom_site. Further enhancements were driven by valuable insights from the PDB data consumer community. Responding to user needs, intra-molecular connectivity for each residue present in an entry was introduced. This addition proved instrumental in encouraging users to transition from the outdated legacy PDB format to the more powerful, PDBx/mmCIF format (https://mmcif.wwpdb.org/dictionaries/mmcif_pdbx_v50.dic/Groups/reference_sequence_group). Researchers can now access the intra-molecular connectivity information by referring to the _chem_comp_bond and _chem_comp_atom categories within the PDBx/mmCIF-formatted files of the PDB NextGen archive (Figure 2). (N.B.: They are also being introduced into the wwPDB FTP archive.) These categories provide comprehensive details on atom connectivity and chemical bonding to support analysis and visualization of the molecules.

**Figure 1.**
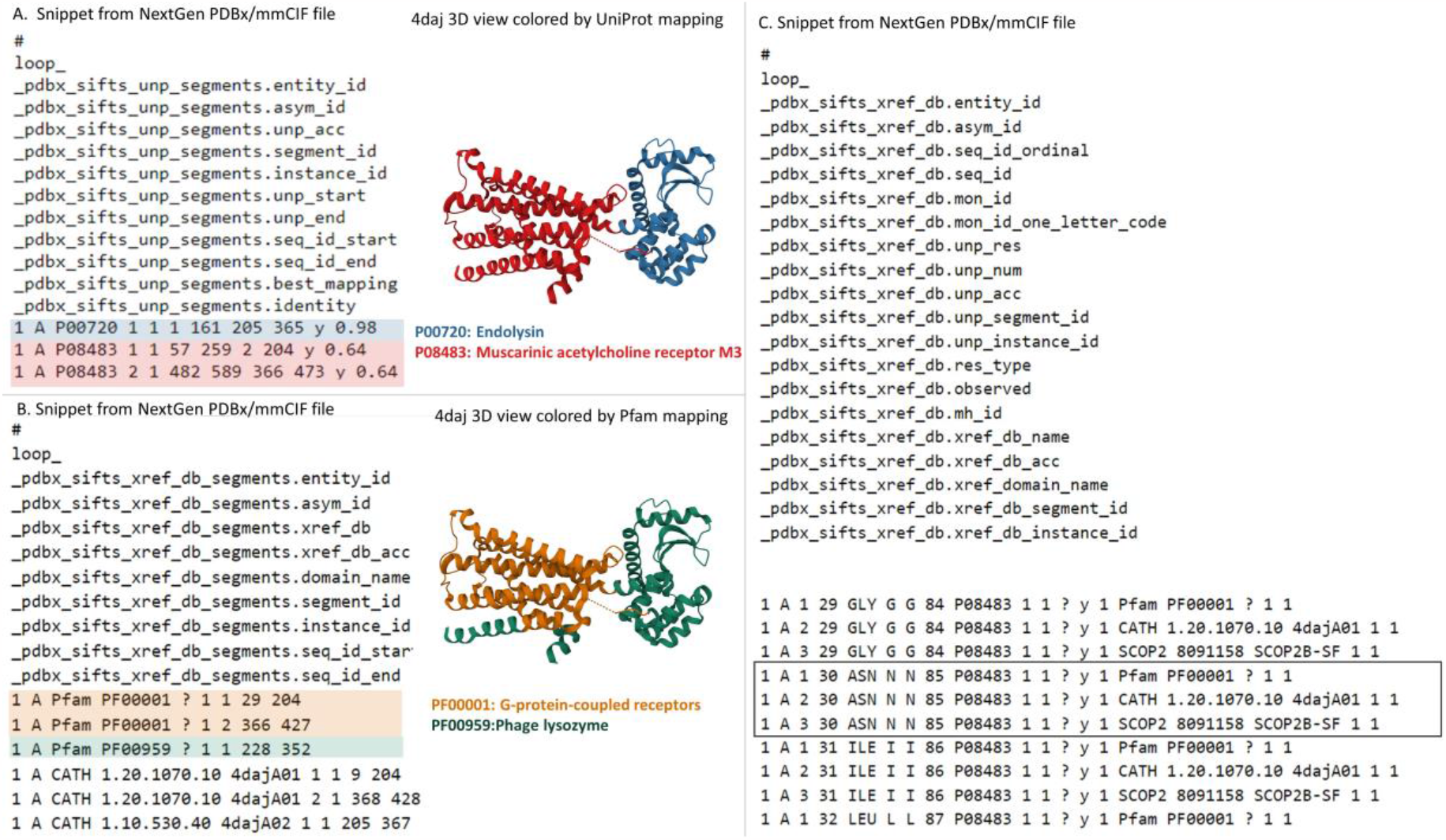
Accessing SIFTS Annotations in the NextGen Archive: This figure displays a snippet from the NextGen Archive PDBx/mmCIF File for PDB ID 4daj, together with a 3D representation of the molecular structure. A) Depicts the “_pdbx_sifts_unp_segments” category, presenting two segments of PDB chain A, each mapped to UniProtKB accessions: P00720 and P08483. This suggests that PDB ID 4daj corresponds to a chimeric protein. B) Illustrates the “_pdbx_sifts_xref_db_segments” category, demonstrating residue range-based cross-references to additional databases like Pfam, SCOP2, and CATH. In this case, PDB chain A is associated with two Pfam domains, corresponding to a G-protein-coupled receptor (Pfam accession: PF00001) and Phage lysozyme (Pfam accession: PF00959). C) Displays the “_pdbx_sifts_xref_db” category, providing a comprehensive view of all mappings for each residue to external databases. Notably, the mappings from UniProt and other cross-reference databases (Pfam/SCOP2/CATH) are highlighted in a box for residue Asn30 in chain A.

**Figure 2.**
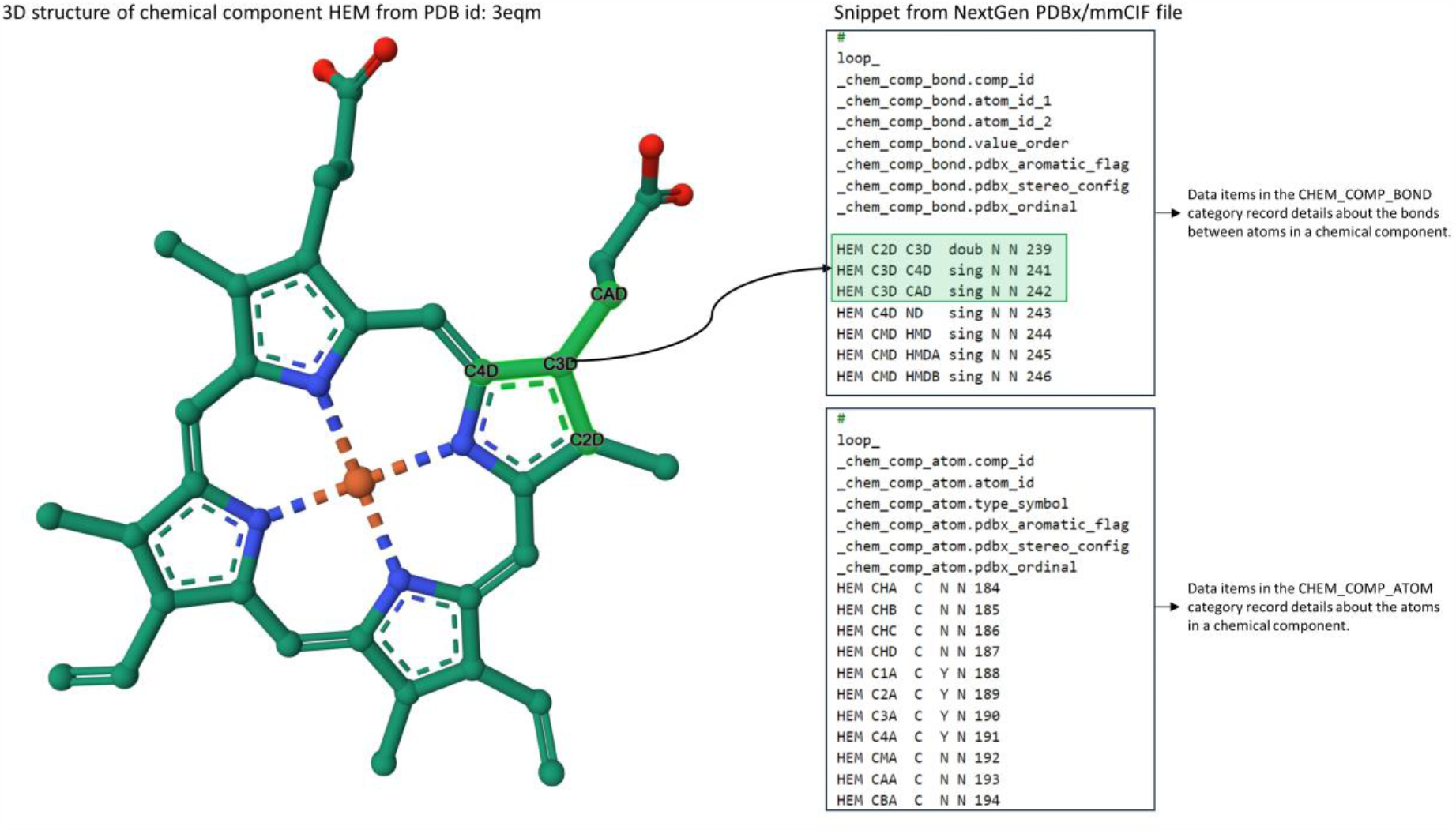
Accessing Intra-molecular Connectivity Information in NextGen Archive: This figure displays a snippet from the NextGen Archive PDBx/mmCIF File and 3D representation of Hemoglobin, identified as the chemical component CCD HEM within PDB ID 3eqm. The “_chem_comp_bond” and “_chem_comp_atom” categories can be used for accessing detailed information about the bonds between atoms within a chemical component and the attributes of individual atoms in that component. Notably, the image highlights a specific instance where atom C3D forms a single bond with atoms C4D and CAD, and a double bond with atom C2D.

New cyber infrastructure was established to collect and combine value-added metadata, utilizing specialized services and the collective expertise of wwPDB partners. To ensure interoperability and data consistency, data enrichment processes at each wwPDB data center generate data that are fully aligned with PDBx/mmCIF semantics. These enrichment processes automatically update evolving annotations wherever this is possible. Doing so not only keeps the data up to date but also minimizes the need for manual intervention, streamlining information dissemination.

Central to this advance was development of a common exchange area, a collaborative data hub wherein each wwPDB partner contributes annotations and metadata. At present, the wwPDB uses the Remote Synchronization (rsync) protocol to exchange data among wwPDB partners. Each wwPDB data center has a private common data exchange area, wherein data can be exchanged *via* rsync. Within the common exchange area, an exhaustive inventory of entries that have undergone updates for each annotation type is maintained. The inventory serves as the foundation for aggregating updated files from each wwPDB partner. As the wwPDB-designated Archive Keeper for the PDB Core Archive, RCSB PDB plays a pivotal role in maintaining all PDB archives (Main, Versioned, and NextGen). A workflow was developed to collate and update the data (Figure 3). Rigorous quality checks are conducted to ensure the validity and accuracy of the merged data files: completeness of SIFTS data, PDBx/mmCIF dictionary compliance within SIFTS data file, and data consistency between SIFTS data and its corresponding PDB data. In case of discrepancies or issues identified during format checking, wwPDB partners are promptly informed, and corrective actions taken. Upon successful merging of these annotations, combined cohesive and enriched PDB data are presented in both PDBx/mmCIF and PDBML formats. Once updated/validated data are ready for distribution, they are synchronised onto a publicly accessible wwPDB NextGen archive, https://files-nextgen.wwpdb.org. This process is fully automated and is currently executed monthly. To enhance accessibility and reduce latency, NextGen archival data are mirrored by RCSB PDB, PDBe, and PDBj.

**Figure 3.**
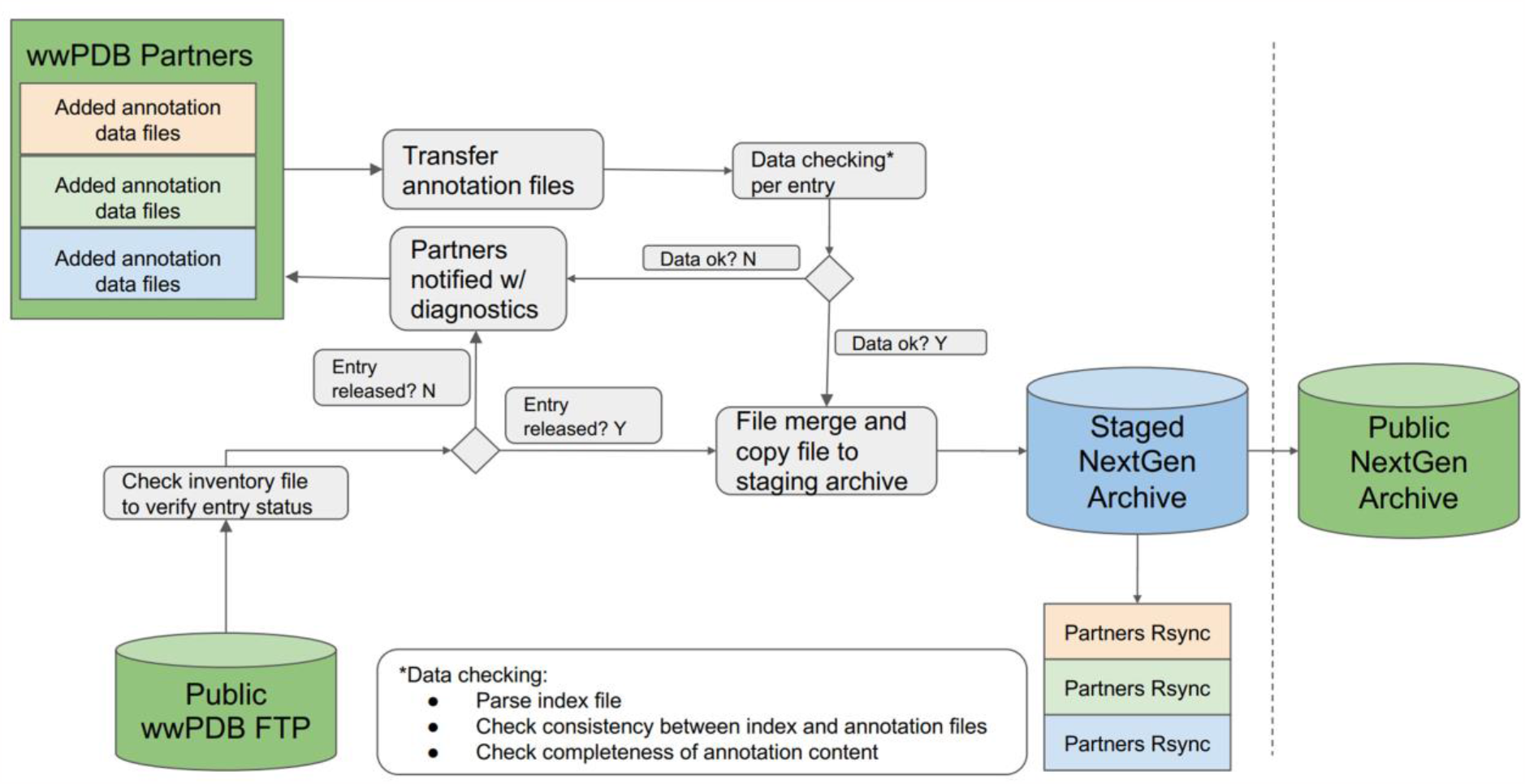
Systematic workflow of NextGen Archive: This figure outlines the structured process for maintaining and updating the NextGen Archive. It showcases key steps, including annotations collection from wwPDB partners, data quality checks, corrective actions, file aggregation and synchronized data in the staging area.

With continuing growth in the size of the PDB archive, issuance of new, longer PDB IDs will become necessary as more and more structures are added (Figure 4). In anticipation of this important milestone in the history of the PDB, the wwPDB partners elected to adopt a revised PDB accession code with a prefix “pdb_” and a length of 8 alphanumeric characters (*e*.*g*., PDB ID 8aly will become pdb_00008aly). This new PDB ID format will offer the added benefits of enabling text mining detection of PDB structures in the scientific literature and allowing more informative and transparent delivery of revised data files. The new extended PDB ID is already being stored in most mmCIF format files under [the_database_2.pdbx_database_accession](https://mmcif.wwpdb.org/dictionaries/mmcif_pdbx_v50.dic/Items/_database_2.pdbx_database_accession.html) data item in addition to the legacy four-character PDB ID, which is stored under _database_2.database_code. Once four-character PDB IDs are fully exhausted, the new extended PDB IDs will be represented in both _database_2.database_code and _database_2.pdbx_database_accession data items. To facilitate a more orderly transition from the legacy PDB format to the PDBx/mmCIF archival standard, NextGen file naming and data now utilize extended PDB IDs. As for the PDB Versioned archive, all data files pertaining to a particular PDB structure in the NextGen archive are stored in a single directory following a 2-character hash from the penultimate two characters of the PDB code, ‘third from last character’ and ‘second from last character’. This hash code will be preserved once PDB IDs are extended to eight characters with the pdb_ prefix. Some examples are provided below:

**Figure 4.**
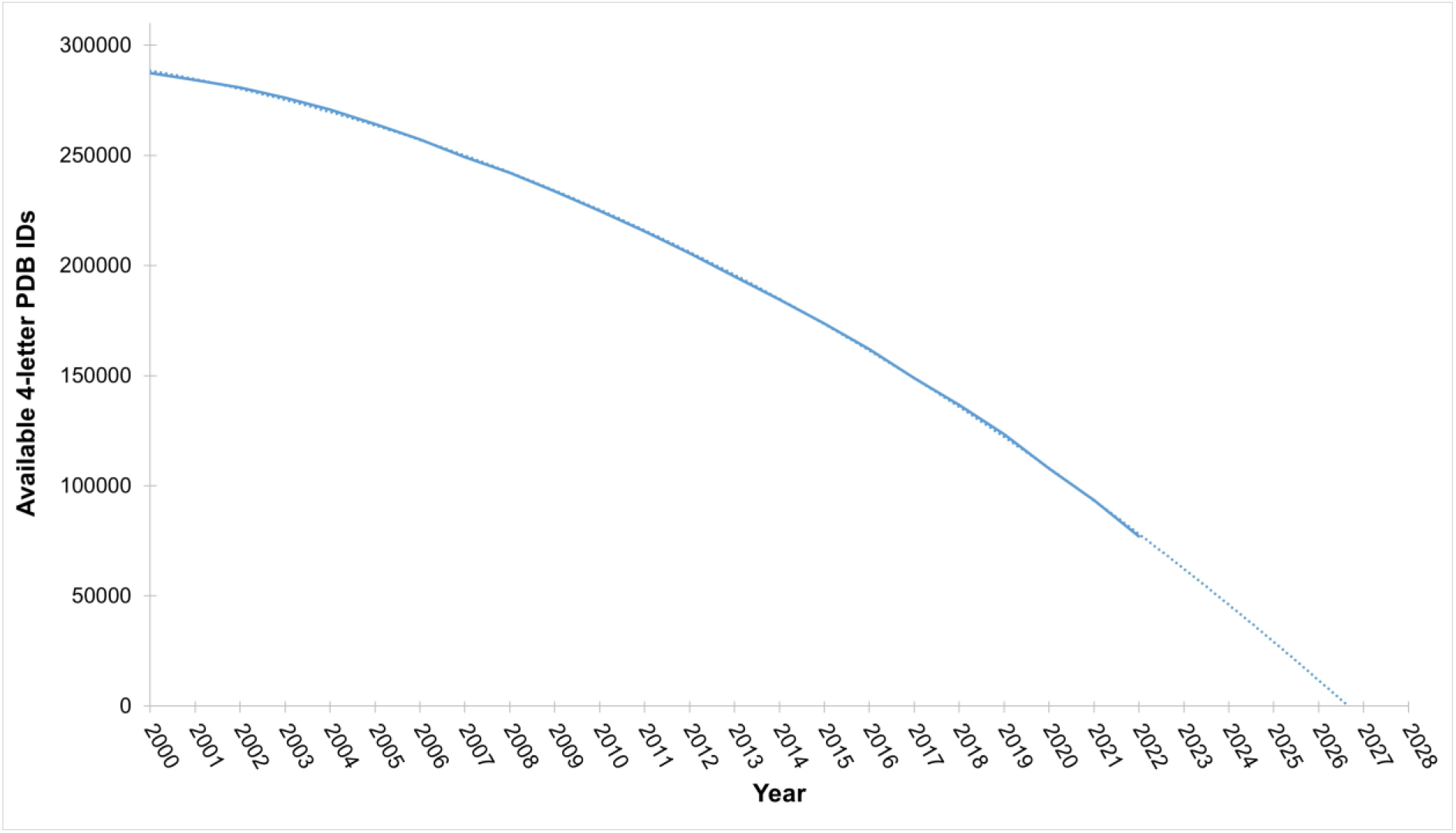
Availability of 4-Letter PDB Codes *versus* Time: This figure depicts the annual count of available 4-letter PDB codes. Current projections anticipate exhaustion of 4-letter PDB codes by the end of 2027.

PDB entry 8aly in the NextGen has PDB identifier pdb_00008aly and is accessible at https://files-nextgen.wwpdb.org/pdb_nextgen/data/entries/divided/al/pdb_00008aly/

Both PDBx/mmCIF and PDBML data files are present at this location (*e*.*g*., for pdb_00008aly: pdb_00008aly_xyz-enrich.cif.gz and pdb_00008aly_xyz-no-atom-enrich.xml.gz are available respectively).

Usage of the PDB NextGen Archive is being tracked continuously to assess breadth of impact. Between February and September 2023, more than 3.5 million enriched NextGen PDB archive data files in PDBx/mmCIF format have been downloaded by PDB data consumers around the world.

## Conclusion

Development of the PDB NextGen Archive cyber infrastructure marks a significant milestone in PDB data integration. Through use of a common exchange area, data enrichment, automated updates, aggregation mechanisms, rigorous quality checks, and global open access, the wwPDB partnership ensures that researchers have access to accurate, up-to-date, and easily accessible structural biology information, facilitating advances and discoveries across diverse scientific disciplines.

With PDBx/mmCIF-based infrastructure at the NextGen Archive, the extensibility of the data model and update of existing files can be achieved easily and independently from the PDB main archive. The plan for the immediate future is to further enrich annotation by providing ligand-binding information for small molecules. wwPDB also plans to expand NextGen Archive content with an investigative study by grouping certain types of entries at the investigation level. The NextGen Archive update schedule is currently monthly.

The wwPDB partnership encourages scientific journals to adopt the new PDB ID format (“pdb_” prefix followed by 8 alphanumeric characters) as soon as possible. Existing entries with 4-character PDB IDs are given new PDB IDs by adding prefixing “pdb_0000” to the four-character IDs when entries are updated (*e*.*g*., the new extended identifier for PDB ID “1abc” is “pdb_00001abc”). Update of existing entries with extended PDB IDs in the _database category will be completed by the end of 2024.

## Availability

The PDB NextGen archive files are accessible *via* wwPDB HTTPS and rsync protocols and its mirrors in the USA, UK, and Japan via the following links respectively:

∘ wwPDB: https://files-nextgen.wwpdb.org, rsync://rsync-nextgen.wwpdb.org
∘ RCSB PDB (USA): https://files-nextgen.rcsb.org, rsync://rsync-nextgen.rcsb.org
∘ PDBe (UK): http://ftp.ebi.ac.uk/pub/databases/pdb_nextgen
∘ PDBj (Japan): https://files-nextgen.pdbj.org, rsync://rsync-nextgen.pdbj.org

At these locations, both PDBx/mmCIF and PDBML format files are provided, with the suffix “_xyz-no-atom-enrich.xml” for PDBML and “_xyz-enrich.cif” for PDBx/mmCIF. More download mechanisms and additional details can be found on the official download page: https://www.wwpdb.org/ftp/pdb-nextgen-archive-site

The PDB NextGen Repository is currently updated on the first Wednesday of each month at 00:00 UTC.

## Funding

Work on the PDB NextGen archive was supported by UK Biotechnology and Biological Research Council (BB/V004247/1, PI: Sameer Velankar) and the NSF (DBI-2019297, PI: S.K. Burley).

## Conflict of interest statement

None declared.

## Notes

http://www.wwpdb.org/ftp/pdb-nextgen-archive-site

### Competing Interest Statement

The authors have declared no competing interest.

